# Variable loss of antibody potency against SARS-CoV-2 B.1.1.529 (Omicron)

**DOI:** 10.1101/2021.12.19.473354

**Authors:** Daniel J. Sheward, Changil Kim, Roy A. Ehling, Alec Pankow, Xaquin Castro Dopico, Darren Martin, Sai Reddy, Joakim Dillner, Gunilla B. Karlsson Hedestam, Jan Albert, Ben Murrell

## Abstract

The recently-emerged SARS-CoV-2 B.1.1.529 variant (Omicron) is spreading rapidly in many countries, with a spike that is highly diverged from the pandemic founder, raising fears that it may evade neutralizing antibody responses. We cloned the Omicron spike from a diagnostic sample which allowed us to rapidly establish an Omicron pseudotyped virus neutralization assay, sharing initial neutralization results only 13 days after the variant was first reported to the WHO, 8 days after receiving the sample.

Here we show that Omicron is substantially resistant to neutralization by several monoclonal antibodies that form part of clinical cocktails. Further, we find neutralizing antibody responses in pooled reference sera sampled shortly after infection or vaccination are substantially less potent against Omicron, with neutralizing antibody titers reduced by up to 45 fold compared to those for the pandemic founder. Similarly, in a cohort of convalescent sera prior to vaccination, neutralization of Omicron was low to undetectable. However, in recent samples from two cohorts from Stockholm, Sweden, antibody responses capable of cross-neutralizing Omicron were prevalent. Sera from infected-then-vaccinated healthcare workers exhibited robust cross-neutralization of Omicron, with an average potency reduction of only 5-fold relative to the pandemic founder variant, and some donors showing no loss at all. A similar pattern was observed in randomly sampled recent blood donors, with an average 7-fold loss of potency. Both cohorts showed substantial between-donor heterogeneity in their ability to neutralize Omicron. Together, these data highlight the extensive but incomplete evasion of neutralizing antibody responses by the Omicron variant, and suggest that increasing the magnitude of neutralizing antibody responses by boosting with unmodified vaccines may suffice to raise titers to levels that are protective.

## Introduction

A new SARS-CoV-2 variant, B.1.1.529 (designated “Omicron” by the WHO) is rapidly replacing the highly transmissible Delta variant (B.1.617.2) in many countries. Relative to the pandemic founder, the archetypical Omicron variant harbors two deletions, one insertion, and 30 amino acid differences in the viral spike, including many known or predicted to confer resistance to neutralizing antibodies^1^. However, their combined effect, and the phenotypic effects of a number of novel Omicron mutations are unknown. Such a substantial antigenic shift may undermine protection afforded by currently licensed vaccines, and monoclonal antibodies used in the clinic. We therefore characterized, using a pseudotyped virus assay, the sensitivity of Omicron to neutralization by relevant monoclonal antibodies, pooled serum from vaccinees, serum samples from infected and infected-then-vaccinated healthcare workers, as well as a random sample of recent seropositive blood donors.

## Results

An Omicron variant spike was molecularly cloned from an anonymized diagnostic sample, suspected to contain B.1.1.529 due to S-gene target failure. A region of spike from before-the-first to after-the-last mutation in Omicron (codons corresponding to amino acid positions 43 to 1000) was PCR amplified from cDNA, subcloned into a spike expression vector, and confirmed by sequencing to encode the Omicron consensus sequence (Supp. Table 1). Using this molecular spike clone, we generated pseudotyped lentiviral particles, and assessed the relative sensitivity of the Omicron variant to neutralization.

**Table 1.**
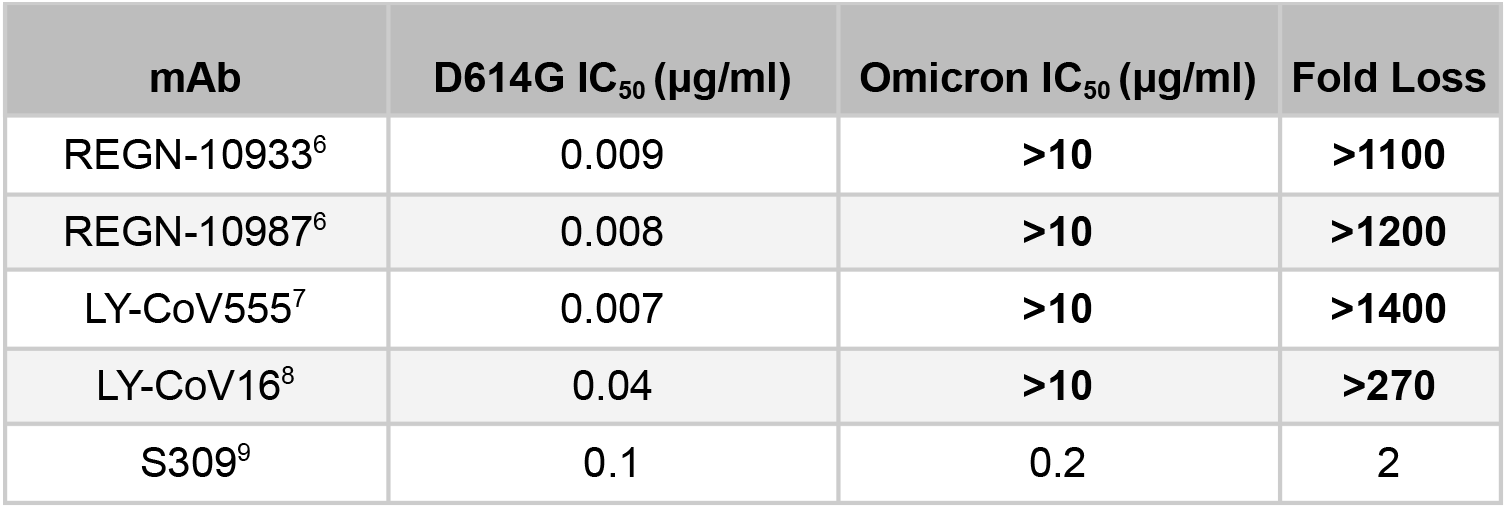
Failure of several monoclonal antibodies included in therapeutic cocktails to neutralize the Omicron variant.

The First WHO International Standard immunoglobulin (20/136) - pooled from convalescent patients in 2020 - showed a ±40-fold reduction in the neutralization of Omicron compared to the pandemic founder (‘wild-type’, WT) (IC_50_ from 0.6 IU/ml to 23.4 IU/ml) (Fig. 1A), indicating substantial resistance to antibodies elicited by ancestral SARS-CoV-2 infection, in line with the significant neutralization resistance observed for Omicron in parallel live virus assays^2^.

**Figure 1.**
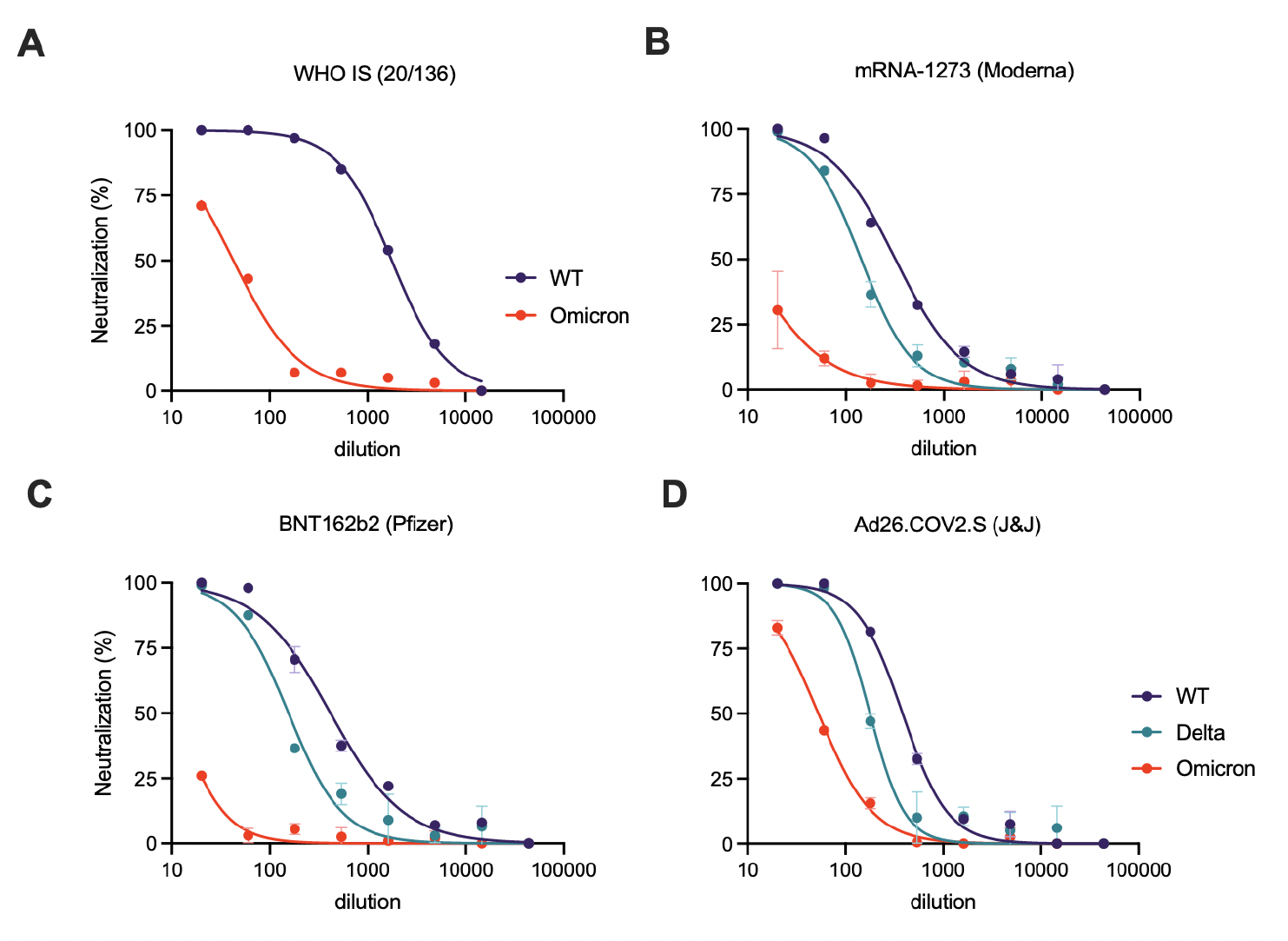
Significant resistance of SARS-CoV-2 B.1.1.529 (Omicron) to antibody neutralization by reference reagents. **A**. Neutralization of the Omicron variant by the First WHO International Standard Immunoglobulin from convalescent individuals. **C-F**. Neutralization of the B.1.1.529 (Omicron) variant compared to wild-type and B.1.617.2 (Delta) by pooled sera standards from vaccinees receiving (**B**) mRNA-1273 (Moderna), (**C**) BNT162b2 (Pfizer/BioNTech), or (**D**) Ad26.Cov2.S (Johnson & Johnson). Error bars depict mean ±SD. The WHO International Standard Immunoglobulin was assayed only once per variant, due to reagent limitations.

Next, to evaluate the likely impact of Omicron on the efficacy of vaccine-elicited antibodies, we assessed the neutralization of Omicron by pooled serum standards from BNT162b2 (Pfizer/BioNTech), mRNA-1273 (Moderna) and Ad26.COV2.S (Johnson&Johnson) vaccine recipients. We found that neutralization of Omicron was substantially reduced, from 7 to 45 fold, across the vaccine standard serum pools (Fig. 1B-D).

While neutralization in serum sampled shortly following vaccination provides critical information about the antibody responses elicited and boosted by vaccines, immunity at the population level and ‘real-world’ vaccine protection incorporates not just vaccination, but a variety of previous and subsequent exposures, as well as the waning^3–5^ of the responses to these. Therefore, to provide a snapshot of status of immunity at the population level, prior to the introduction of Omicron, we assessed neutralization by sera from two cohorts from Stockholm, Sweden: (i) 17 randomly selected seropositive recent blood donors (anonymized and therefore unknown exposure and vaccination status) and (ii) 17 recently-sampled hospital workers (HW) who were infected in May 2020, with varied subsequent vaccination histories (Supp. Table 2).

Neutralizing ID_50_ titers for the blood donors were ±7-fold lower against Omicron compared to WT (Fig 2B). However, the reduction in neutralizing activity was heterogeneous, with some sera nearly 25-fold less potent and others experiencing no significant loss, indicating the presence of cross-neutralizing antibodies in a subset of donors. Similarly, sera from hospital workers were, on average, ±5-fold less potent against Omicron and also exhibited considerable inter-individual variation (Fig. 2B).

**Figure 2:**
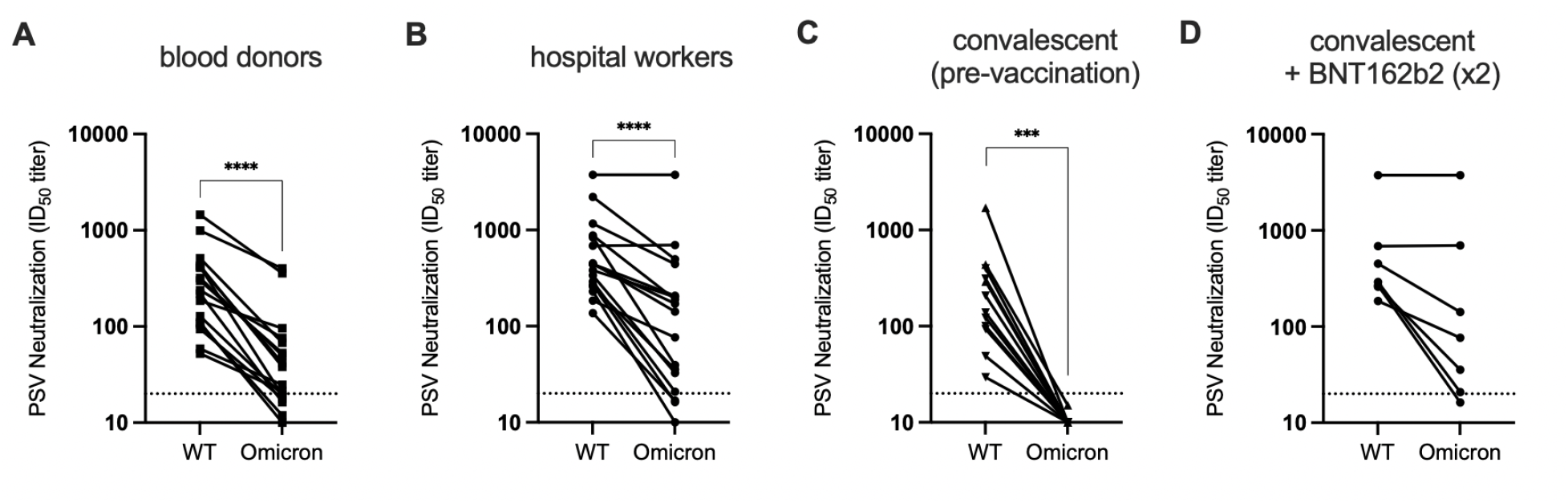
Characterization of the relative neutralization resistance of Omicron. **A-B**. Paired WT and Omicron pseudovirus neutralization titers for the Recent Blood Donor (BD; N=17) and Hospital Worker (HW; N=17) cohorts against the pandemic founder (WT) and B.1.1.529 (Omicron). **C**. Samples taken from the HWs approximately one (▲) or four (▼) months after infection, but prior to vaccination (convalescent) show a substantial reduction in the ability to neutralize Omicron, with all ID_50_s falling below the limit of detection (20). **D**. For seven HW that reported receiving two doses of BNT162b2 subsequent to infection, cross-neutralization of Omicron was evident.

Importantly, historical samples taken from the HW cohort after confirmed SARS-CoV-2 infection, but prior to vaccination (convalescent), showed a near-complete loss of neutralizing activity against Omicron (Fig. 2C). However, for seven hospital workers that received two doses of BNT162b2 after infection, robust cross-neutralization of Omicron was evident in a number of individuals (Fig. 2D), highlighting the improvement in the neutralisation of variants afforded by vaccination in previously infected individuals.

Monoclonal antibodies represent important treatment and prophylactic options for certain categories of patients, and can significantly reduce morbidity in those otherwise at risk for severe COVID-19^10^. We therefore evaluated the sensitivity of the Omicron variant to neutralization by several monoclonal antibodies currently included in therapeutic cocktails used in the clinic. REGN10933, REGN10987, Ly-CoV016 and Ly-CoV555, all failed to neutralize Omicron up to the highest concentration tested (10 μg/ml) despite potently neutralizing the ancestral B.1 (D614G) spike (Table 1). However, the parent of Sotrovimab, S309, maintained much of it’s activity, experiencing only a 2-fold loss in potency against Omicron (Table 1).

## Discussion

Neutralizing antibodies are a mechanistic correlate of SARS-CoV-2 vaccine protection^11,12^. While other arms of the immune system contribute to protection from severe disease, the significant reduction in neutralization sensitivity that we document here will likely translate into an erosion of vaccine-mediated protection. This is supported by the recent, rapid spread of Omicron in countries with high vaccine coverage^13^, as well as preliminary reports of multiple breakthrough infections^14^.

We show that there is a precipitous drop in neutralization potency against Omicron for serum pools from convalescent donors and recently vaccinated individuals, as well as from individual convalescent donors sampled soon after initial infection. However, sera from a high-risk cohort^15^ of infected-then-vaccinated health care workers exhibit substantial cross-neutralization of Omicron, which correlates with their ability to cross-neutralize other variants. This suggests that responses to SARS-CoV-2 spike broaden with increasing antigenic exposure. This has been characterized against other variants in the context of both prior infections^16^ and three-dose vaccinations^17^.

Interestingly, responses in a cohort of random recently-sampled Stockholm seropositive blood donors also exhibited substantial cross-neutralization of Omicron. On average, the fold loss against Omicron was only slightly greater than that of the previously infected hospital worker cohort. Such cross neutralization in a “real world” cohort would not have been predicted from the responses observed at the peak post-vaccination^2^. As most individuals in Sweden have only received two vaccine doses, this breadth may then be explained by the frequency of exposures prior to, or following, vaccination in Stockholm, Sweden. Alternatively, there may be systematic differences in cross neutralization for samples taken immediately after a second immunization compared to those sampled later. Indeed, affinity maturation of antibody lineages over the course of months after SARS-CoV-2 infection enabled the cross-neutralization of variants of concern, and heterologous sarbecoviruses^18^.

From a global health perspective, the dramatic loss of neutralization against Omicron for previously-infected but unvaccinated individuals has implications for whether such individuals can be considered immune. Further, the cross-neutralizing antibody responses in the infected-then-vaccinated hospital worker cohort indicate significant value in vaccinating the previously-infected.

Given the complete resistance of Omicron to several monoclonal antibodies currently included in cocktails used in the clinic, treatment options should be informed by rapid SARS-CoV-2 genotyping in regions where Omicron and Delta (or other variants) are both circulating. This also argues for the rapid diversification of our clinical monoclonal antibody portfolio, to hedge against unpredictable potency losses for future variants. It also highlights the need to rapidly screen variants for their sensitivity to clinical therapeutics.

Methodologically, the current standard practice for generating pseudovirus spike expression plasmids for novel variants relies on site-directed mutagenesis when only a small number of mutations differ from an existing plasmid construct, or gene synthesis to generate entire spike genes. Exceptional urgency is demanded by the emergence of a rapidly-spreading novel variant with a large number of spike mutations. Molecular cloning from a diagnostic sample allowed us to circumvent gene synthesis delays, and share pseudovirus neutralization data just eight days after receipt of the suspected Omicron diagnostic samples, and 13 days after the variant was first reported to the WHO^19^. One risk associated with this approach is that, if the expression of a non-codon optimized spike is too low, pseudovirus entry into target cells may be too inefficient to accurately quantify neutralization. For this reason, our cloning strategy retained as much of the codon-optimized backbone as possible, especially in the C-terminal region of the spike, which is not mutated in Omicron. It is not clear whether such a strategy would universally succeed with all variants, so a dual approach that attempts gene synthesis and direct cloning (when samples are available) would mitigate this risk.

Ultimately, long-term protection against SARS-CoV-2, including antigenic variants that will arise, may require updated vaccines or vaccines that elicit more broadly cross-neutralizing antibodies. Until such vaccines are available, our data from two different cohorts suggests that the loss of neutralization against Omicron is incomplete. It has previously been shown, with other variants, that a third dose with unmodified vaccines may have a broadening effect^17^. But even without such a booster broadening effect, in many donors the magnitude of loss in neutralization we observe against Omicron argues that antibody titers may be boostable into a protective range with currently licensed vaccines.

## Acknowledgements

We acknowledge the G2P-UK National Virology consortium funded by MRC/UKRI (grant ref: MR/W005611/1) and the Barclay Lab at Imperial College for providing B.1, B.1.351, B.1.617.2, and B.1.621 spike-encoding plasmids. We acknowledge Penny Moore and the NICD (South Africa) for providing a B.1.351 spike plasmid, which was generated using funding from the South African Medical Research Council. pCMV-dR8.2 dvpr was a gift from Bob Weinberg (Addgene plasmid # 8455; http://n2t.net/addgene:8455; RRID:Addgene_8455). pBOBI-FLuc was a gift from David Nemazee (Addgene plasmid # 170674; http://n2t.net/addgene:170674; RRID:Addgene_170674). We acknowledge all staff at the Department of Clinical Microbiology, Karolinska University Hospital involved in SARS-CoV-2 routine diagnostics, S-gene screening and sequencing.

The following reagents were obtained through NIAID, NIH:

Pooled Human Serum Sample, Pfizer Vaccine (BEI Resources: NRH-17727)

Pooled Human Serum Sample, Moderna Vaccine (BEI Resources: NRH-17846)

Pooled Human Serum Sample, Johnson&Johnson Vaccine (details unavailable at the time of submission).

This project was supported, in part, by funding from the European Union’s Horizon 2020 research and innovation programme under grant agreement no. 101003653 (CoroNAb) to G.B.K.H., S.R., and B.M; from the SciLifeLab Call 4.1: Laboratory preparedness for pandemics (Reg no. VC-2021-0033) to B.M. and J.A.; from the Erling Persson Foundation to B.M and G.B.K.H.

## Author Contributions

Conceptualization, D.J.S., D.M., G.B.K.H., J.A., B.M.; Formal analysis, D.J.S., B.M.; Investigation, D.J.S., C.K., X.C.D; Methodology, A.P., D.J.S., B.M.; Visualization, D.J.S., B.M.; Resources, R.E., S.R, J.D., G.B.K.H., J.A.; Supervision, D.J.S., G.B.K.H., S.R, J.A., B.M; Writing – original draft, D.J.S. and B.M.; Writing – review & editing, D.J.S, D.M., G.B.K.H, J.A., B.M.

## Methods

### Ethics statement

HW and Convalescent cohorts: Informed consent was obtained from all participants as part of an ethics approval (Decision number 2020-01620, with amendments 2020-02881 and 2020-05630) from the Swedish Ethical Review Authority. BD cohort and the Omicron-positive sample from which the spike was cloned were anonymized, and not subject to ethical approvals, as per advisory statement 2020–01807 from the Swedish Ethical Review Authority.

### Donor sample description

Two cohorts were studied. Cohort 1 comprised serum samples with detectable neutralization against the Wu-Hu-1 founder variant from 17 anonymized blood donors (“BD”), donated during week 48, 2021, in Stockholm, Sweden. Cohort 2 comprised 17 serum samples from Hospital Workers (“HW”) at the Karolinska University Hospital in Stockholm^20^, who were invited to participate in a study that aimed to characterize their antibody responses following SARS-CoV-2 infection and subsequent vaccinations. Participants, confirmed PCR positive, had serum sampled in April/May 2020, in June/July 2020 (“convalescent”, prior to vaccination), and again in November 2021 (“HW”). Convalescent samples are from 9 unique donors, with 3 donors sampled in both April/May 2020 and June/July 2020. Statistical comparison was performed on the 9 samples from June/July 2020 only, to avoid non-independence due to repeated sampling of 3 donors.

### Spike expression plasmids

Spike plasmids encoding the B.1, B.1.621, and B.1.617.2 were kindly provided by the G2P-UK National Virology consortium funded by MRC/UKRI (grant ref: MR/W005611/1.) and the Barclay Lab at Imperial College.

An Omicron variant spike was molecularly cloned from an anonymized diagnostic sample, suspected to contain B.1.1.529 due to S-gene target failure and subsequently confirmed by sequencing. A region of spike (with codons corresponding to amino acid positions 43 to 1000) incorporating all of the Omicron variant reference mutations was amplified from cDNA derived from a later-confirmed B.1.1.529 clinical sample obtained from a set of anonymized early cases of suspected Omicron infections. A first PCR round amplified the entire spike gene, and then Gibson assembly overhangs were introduced with a second-round PCR, exploiting regions of existing homology between the codon-optimized parent plasmid and the codon-native spike, in order to maximize overhang length while still keeping primer length under 35bp, allowing for overnight primer synthesis. The N-terminal and C-terminal flanking regions of the parent plasmid were similarly amplified.

Primers (5’-3’) used for the construction of the Omicron Spike expression plasmid

#### Spike PCR

##### First round (primers from ^21^)

SARSCoV1200_22_LEFT GTGATGTTCTTGTTAACAACTAAACGAACA

SARSCoV1200_24_RIGHT ATGAGGTGCTGACTGAGGGAAG

##### Second round

FWD_N_term_sample CAAGGTGTTCAGATCCTCAGTTTTACATTCAACTC

REV_C_term_sample TCTGCAGTCTGCCTGTGATCAACCTATCAATTTGC

#### N-terminus flank PCR

Fwd_CMV_plasmid ACGCAAATGGGCGGTAGGCGTG

REV_N_term_plasmid TGAATGTAAAACTGAGGATCTGAACACCTTGTCGG

#### C-terminus flank PCR

FWD_C_term_plasmid AAATTGATAGGTTGATCACAGGCAGACTGCAGAGC

Rev_plasmid TGGCAACTAGAAGGCACAGTCGAG

The three PCR products were cloned by Gibson Assembly into a restriction-enzyme (NheI and XbaI) digested, codon-optimized SARS-CoV-2 Spike expression vector (in pcDNA3.1) harbouring a mutation that introduces a stop codon that truncates the last 19 amino acids of the cytoplasmic tail (facilitating efficient incorporation onto lentiviral particles). The resulting spike-encoding expression vector was confirmed by sequencing to encode an amino acid sequence identical to that of the Omicron consensus.

### Cloned Omicron Spike coding sequence (with 19AA CT truncation)

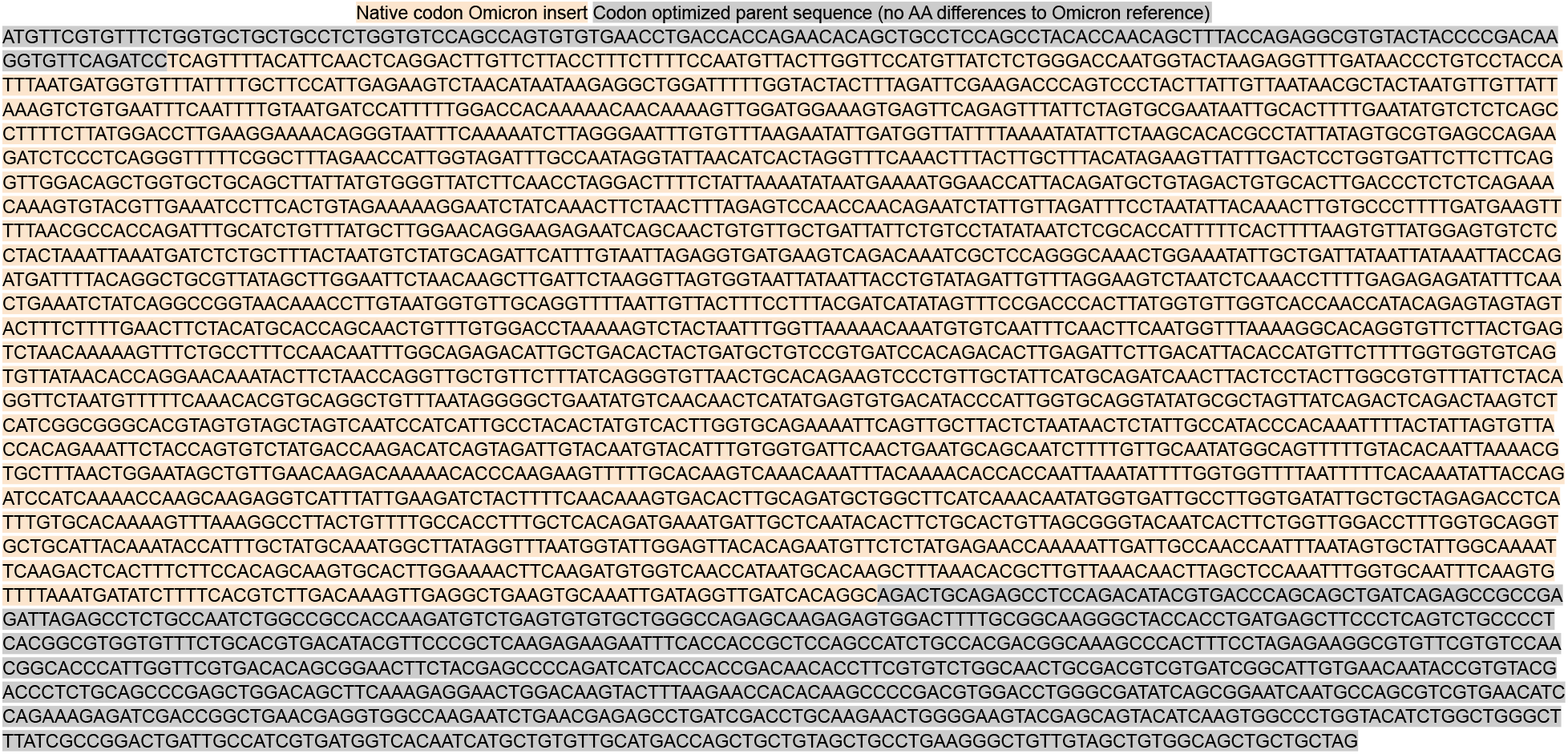

### Cell culture

HEK293T cells (ATCC CRL-3216) and HEK293T-ACE2 cells (stably expressing human ACE2) were cultured in Dulbecco’s Modified Eagle Medium (high glucose, with sodium pyruvate) supplemented with 10% fetal calf serum, 100 units/ml Penicillin, and 100 μg/ml Streptomycin. Cultures were maintained in a humidified 37°C incubator (5% CO_2_).

### Pseudovirus Neutralization Assay

Pseudovirus neutralization assay was performed as previously^22^. Spike-pseudotyped lentivirus particles were generated by the co-transfection of HEK293T cells with a relevant spike plasmid, an HIV gag-pol packaging plasmid (Addgene #8455), and a lentiviral transfer plasmid encoding firefly luciferase (Addgene #170674) using polyethylenimine (PEI).

Neutralization was assessed in HEK293T-ACE2 cells. Briefly pseudoviruses sufficient to produce ±30,000 RLU were incubated with serial 3-fold dilutions of serum for 60 minutes at 37°C in a black-walled 96-well plate. 10,000 HEK293T-ACE2 cells were then added to each well, and plates were incubated for 48 hours. Luminescence was measured using Bright-Glo (Promega) on a GloMax Navigator Luminometer (Promega). Neutralization was calculated relative to the average of 8 control wells infected in the absence of serum. All fold-change comparisons used ID_50_ values from neutralization assays run side-by-side.

### Monoclonal antibody production

Antibody sequences were extracted from deposited RCSB entries and codon optimized (using a human germline-aware codon optimization strategy), then synthesized as gene fragments and cloned into pTWIST transient expression vectors by Gibson assembly or restriction cloning (NotI, BamHI). Cloned plasmids were verified by sanger sequencing.

Expi293 cells were transfected with plasmids encoding for the heavy and light chain at 1µg/mL (i.e. 0.5µg/mL each) according to the manufacturer’s instructions (ThermoFisher, manual for Cat#A14525). Dense Expi293 cultures (day 5-7 post transfection) were centrifuged at 300xg for 5 minutes to pellet the cells. Supernatant was filtered using Steriflip® 0.22 µm (Merck, SCGP00525) filter units. Expi supernatant was directly loaded onto Protein G Agarose (Pierce, Cat# 20399) gravity columns, washed twice with PBS and eluted using Protein G Elution Buffer (Pierce, Cat# 21004). The eluted fractions were immediately neutralized with 1M TRIS-Buffer (pH = 8) to physiological pH. Absorption at 280 nm was quantified by Nanodrop™ 2000c to determine protein containing fractions. These fractions were then pooled and buffer exchanged using SnakeSkin™ dialysis tubing (10 MWCO, Pierce Cat#68100) followed by further dialysis and concentration using Amicon Ultra-4 10kDa centrifugal units (Merck, Cat# UFC801096). The clinically relevant mAbs tested here were in-house produced versions of: REGN-10933^6^, REGN-10987^6^, LY-CoV555^7^, LY-CoV016^8^, and S309^9^ (from which Sotrovimab was derived through Fc modifications).

### Statistical analysis

Individual ID_50_ values for each sample against each variant were calculated in Prism v9 (GraphPad Software) by fitting a four-parameter logistic curve, to neutralization by serial 3-fold dilutions of serum. Comparisons of titers across variants used non-parametric Wilcoxon matched-pairs tests. P values are summarized as: ns P>0.05; * P<0.05; ** P<0.01; *** P<0.001; **** P<0.0001.

## Supplementary Information

**Supplementary Figure 1.**
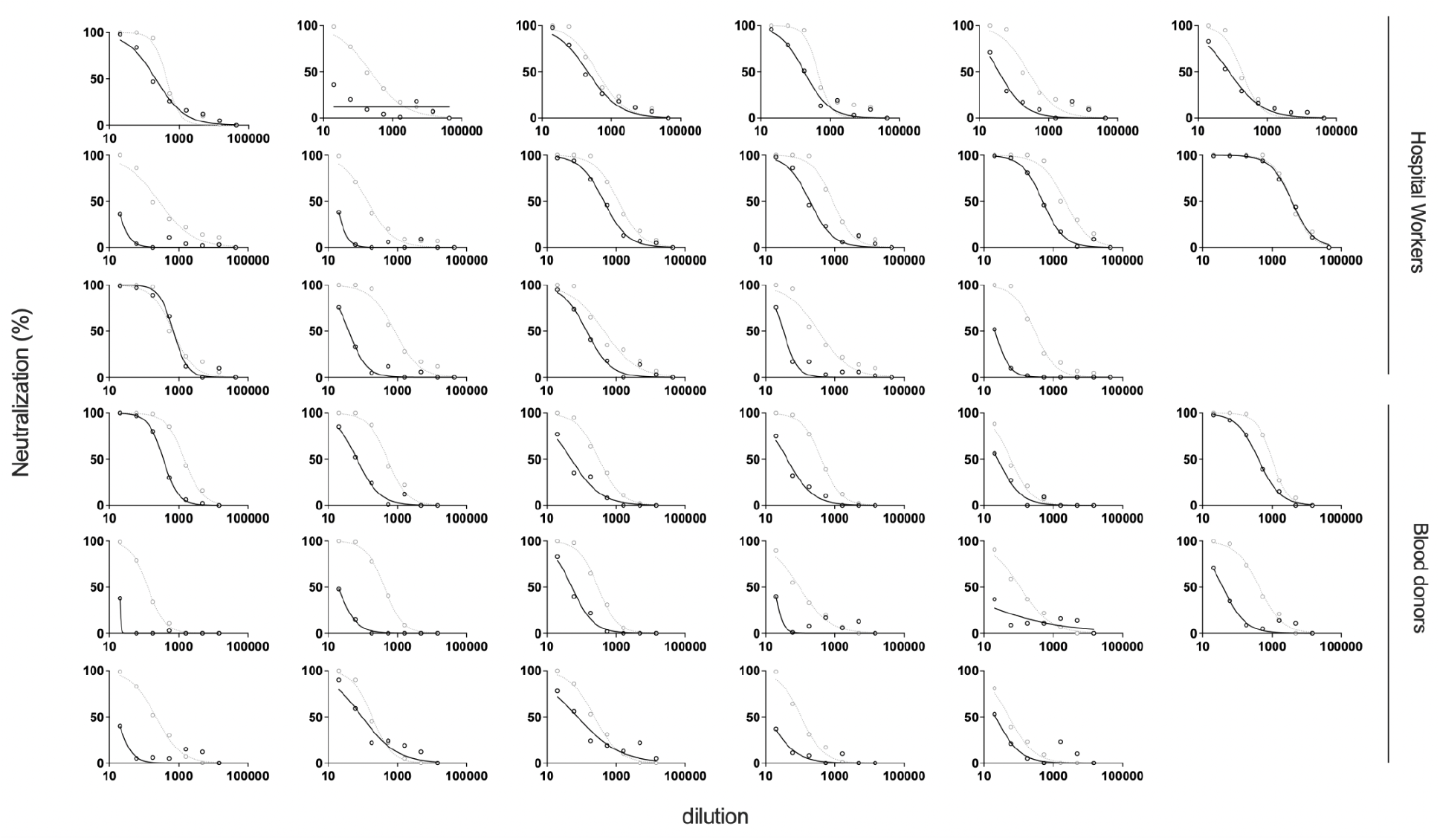
Neutralization curves for each serum from the HW and BD cohorts against WT (grey) and Omicron/B.1.1.529 (black).

**Supplementary Figure 2.**
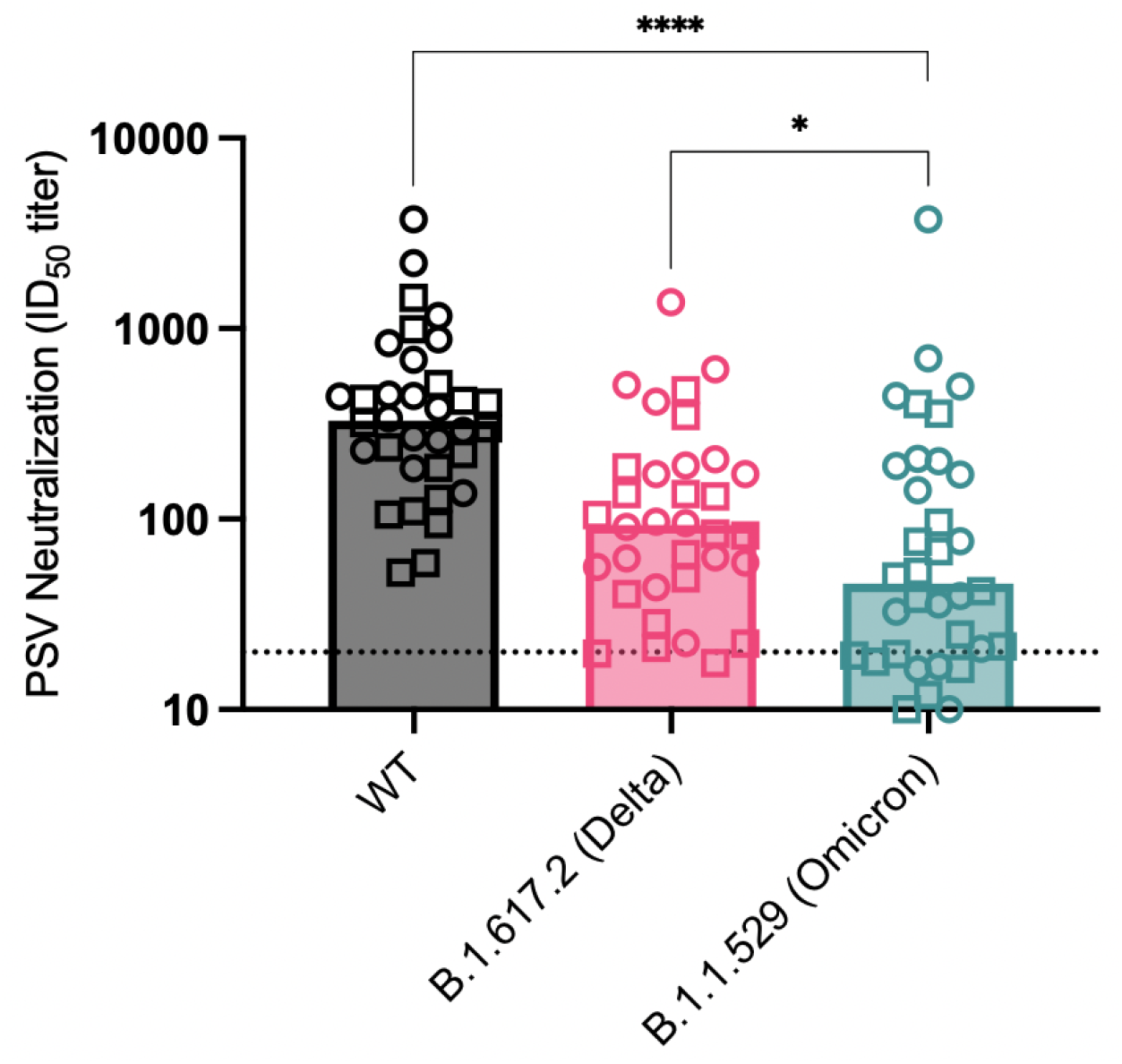
Neutralizing activity of sera from blood donors (BD) (squares) and Hospital Workers (HW) (circles) against WT, Delta, and Omicron.

**Supplementary Figure 3.**
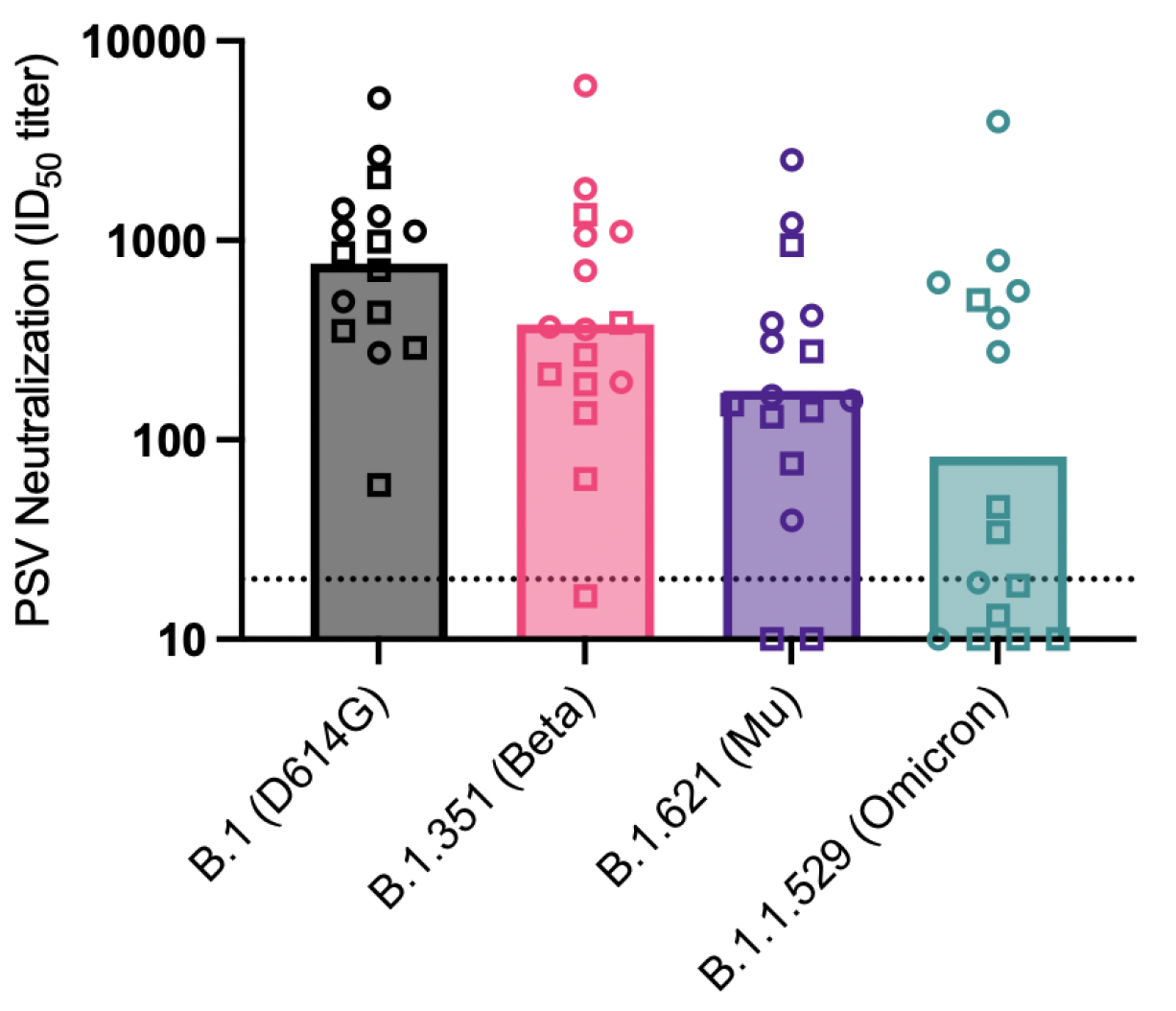
Neutralizing activity of sera from a subset of BD (squares) and HW (circles) against B.1, Beta, Mu, and Omicron, from an independent assay replicate. Sample selection was biased towards samples that showed extreme maintenance or loss against Omicron. For the subset of samples included in both runs, Omicron titers were reproducible (rho = 0.96, calculated from IC50s in the log domain).

**Supplementary Figure 4.**
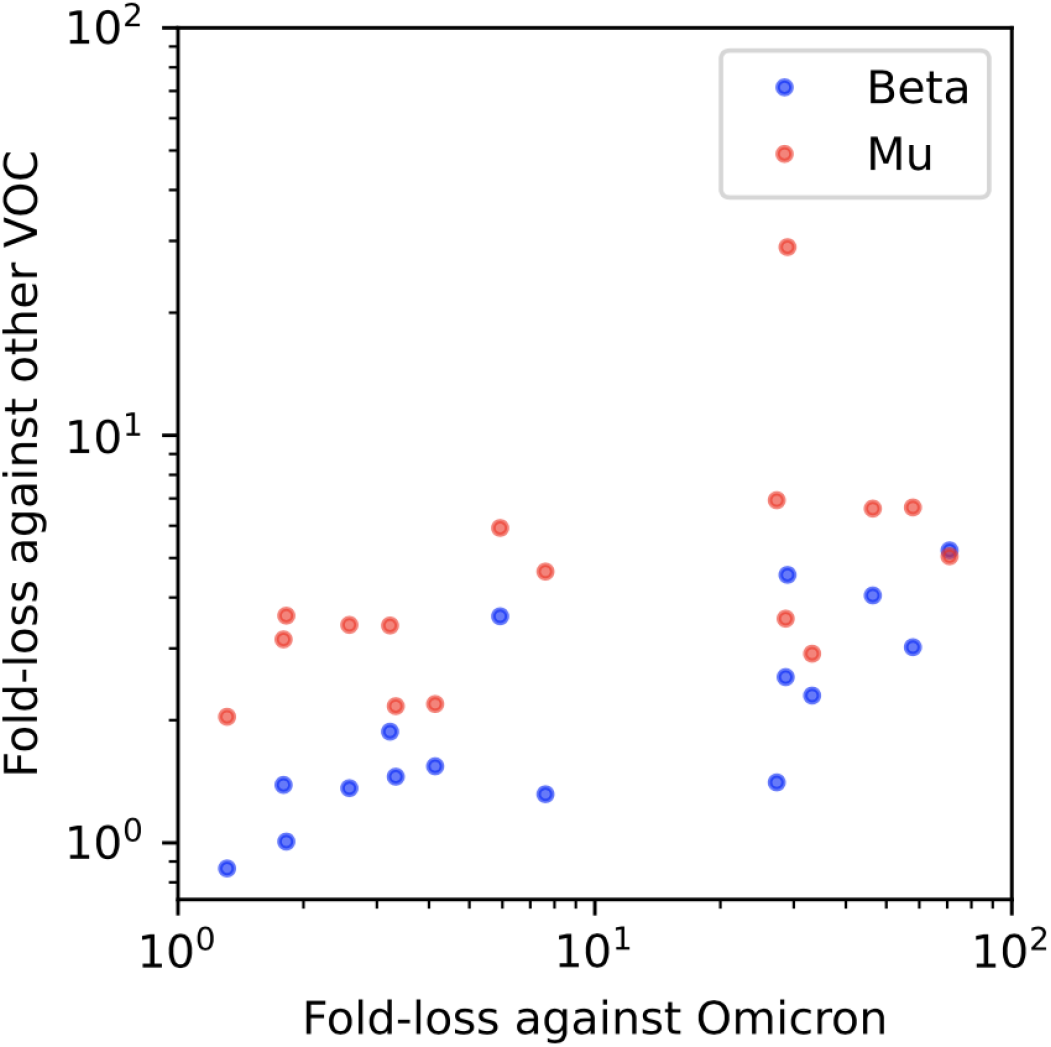
For the subset of samples run against D614G, Omicron, Beta, and Mu, the fold-loss against Omicron (defined as D614G IC50/Omicron IC50) was more strongly correlated with fold-loss against Beta (rho = 0.78, log domain) than Mu (rho = 0.57, log domain).

**Supplementary Table 1.**
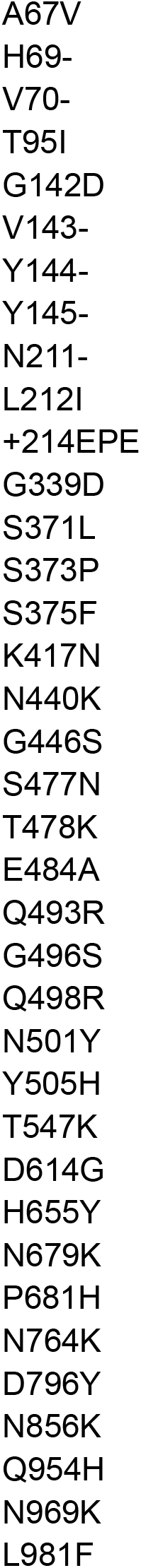
List of mutations relative to Wu-Hu-1 in the Omicron spike evaluated here:

**Supplementary Table 2.** Self-reported vaccination histories of Hospital Workers subsequent to previous infection in early 2020.

| ID | Founder IC50 | Omicron IC50 | Fold reduction | Self reported vaccine history <sup>a</sup> |
| --- | --- | --- | --- | --- |
| A | 839.9 | 42.84 | 19.61 | Unknown |
| B | 442.1 | 213 | 2.08 | AZ, Pfizer (4m) |
| C | 686.3 | 734.4 | 0.93 | Pfizer, Pfizer (4m) |
| D | 184.8 | 78.71 | 2.35 | Pfizer, Pfizer (8m) |
| E | 879.3 | 214.1 | 4.11 | AZ, AZ (5m) |
| F | 230.1 | 10 | 23.01 | Unknown |
| G | 136.6 | 16.69 | 8.18 | AZ, Pfizer (4m) |
| H | 259.4 | 36.39 | 7.13 | Pfizer, Pfizer (8m) |
| I | 439.5 | 179.1 | 2.45 | AZ, AZ (4m) |
| J | 2202 | 549.3 | 4.01 | Unknown |
| K | 381.8 | 207.1 | 1.84 | AZ, AZ (5m) |
| L | 451 | 155.2 | 2.91 | Pfizer , Pfizer, (5m) |
| M | 338.2 | 35.85 | 9.43 | Unknown |
| N | 269 | 15.62 | 17.22 | Pfizer, Pfizer (8m) |
| O | 3748 | 4053 | 0.92 | Pfizer, Pfizer (2m) |
| P | 291.4 | 21.81 | 13.36 | Pfizer, Pfizer (4m) |
| Q | 1166 | 492.7 | 2.37 | Unknown |

<sup>a</sup>Calendar months between most recent immunization and HW serum sampling day shown in brackets.

